# ATM-dependent formation of a novel chromatin compartment regulates the Response to DNA Double Strand Breaks and the biogenesis of translocations

**DOI:** 10.1101/2021.11.07.467654

**Authors:** Coline Arnould, Vincent Rocher, Aldo S. Bader, Emma Lesage, Nadine Puget, Thomas Clouaire, Raphael Mourad, Daan Noordermeer, Martin Bushell, Gaëlle Legube

**Affiliations:** MCD, Centre de Biologie Intégrative (CBI), CNRS, Université de Toulouse, UT3; Cancer Research UK Beatson Institute, Garscube Estate, Switchback Road, Bearsden, Glasgow G61 1BD, UK; Université Paris-Saclay, CEA, CNRS, Institute for Integrative Biology of the Cell (I2BC), 91198, Gif-sur-Yvette, France; Institute of Cancer Sciences, University of Glasgow, Garscube Estate, Switchback Road, Bearsden, Glasgow G61 1QH, UK

## Abstract

DNA Double-Strand Breaks (DSBs) repair is essential to safeguard genome integrity but the contribution of chromosome folding into this process remains elusive. Here we unveiled basic principles of chromosome dynamics upon DSBs in mammalian cells, controlled by key kinases from the DNA Damage Response. We report that ATM is responsible for the reinforcement of topologically associating domains (TAD) that experience a DSB. ATM further drives the formation of a new chromatin sub-compartment (“D” compartment) upon clustering of damaged TADs decorated with γH2AX and 53BP1. “D” compartment formation mostly occurs in G1, is independent of cohesin and is enhanced upon DNA-PK pharmacological inhibition. Importantly, a subset of DNA damage responsive genes that are upregulated following DSBs also physically localize in the D sub-compartment and this ensures their optimal activation, providing a function for DSB clustering in activating the DNA Damage Response. However, these DSB-induced changes in genome organization also come at the expense of an increased translocations rate, which we could also detect on cancer genomes. Overall, our work provides a function for DSB-induced compartmentalization in orchestrating the DNA Damage Response and highlights the critical impact of chromosome architecture in genomic instability.

## Main

DNA Double-Strand Breaks (DSBs) are highly toxic lesions that can trigger translocations or gross chromosomal rearrangements, thereby severely challenging genome integrity and cell homeostasis. Chromatin plays a pivotal function during DNA repair, which is achieved by either non-homologous end joining or homologous recombination pathways^1^. Yet, little is known about the contribution of chromosome architecture into these processes. DSBs activate the DNA Damage Response (DDR) that largely relies on PI3K kinases, including ATM and DNA-PK, and on the establishment of megabase-sized, γH2AX-decorated chromatin domains that act as seeds for subsequent signaling events, such as 53BP1 recruitment and DDR foci formation^2,3^.

Importantly, γH2AX spreading is largely influenced by the pre-existing chromosome conformation in topologically associating domains (TADs)^4–6^ and we recently reported that loop-extrusion, which compacts the chromatin and leads to TADs formation, is instrumental for γH2AX spreading and DDR foci assembly^5^. Moreover, irradiation induces a general chromatin response reinforcing TADs genome wide^7^. At a larger scale, previous work in mammalian cells revealed that DSBs display the ability to “cluster” within the nuclear space (*i*.*e*., fuse) forming large microscopically visible repair foci, composed of several individual repair foci^8–10^. DSB clustering depends on the actin network, the LINC (a nuclear envelope embedded complex)^9,11,12^, as well as on the liquid-liquid phase separation properties of 53BP1^13,14^. The function of DSB clustering has remained enigmatic given that juxtaposition of several DSBs can elicit translocation (*i*.*e*: illegitimate rejoining of two DNA ends)^10^, questioning the selective advantage of DSB clustering/ repair foci fusion^15^.

### ATM drives an acute reinforcement of damaged TADs

In order to get comprehensive insights into chromosome behavior following DSBs, we analyzed 3D genome organization using Hi-C data generated in the human DIvA cell line where multiple DSBs are induced at annotated positions upon hydroxytamoxifen (OHT) addition^16^. Our previous analyses using γH2AX ChIP-seq and direct DSB mapping by BLESS allowed us to identify 80 robustly induced DSBs on the human genome^3^. Using differential Hi-C maps, we found that intra-TAD contacts frequencies were strongly increased within TADs that experience a DSB (*i*.*e*. damaged TADs, Fig. 1a, right panel red square) compared to undamaged TADs, while contacts with neighboring adjacent domains were significantly decreased (Fig. 1a, right panel blue square, Fig. 1b). Interestingly, in some instances, the DSB itself displayed a particularly strong depletion of contact frequency with adjacent chromatin (Fig. 1c black arrow) indicating that the DSB is kept isolated from the surrounding environment, outside of its own TAD.

**Figure 1:**
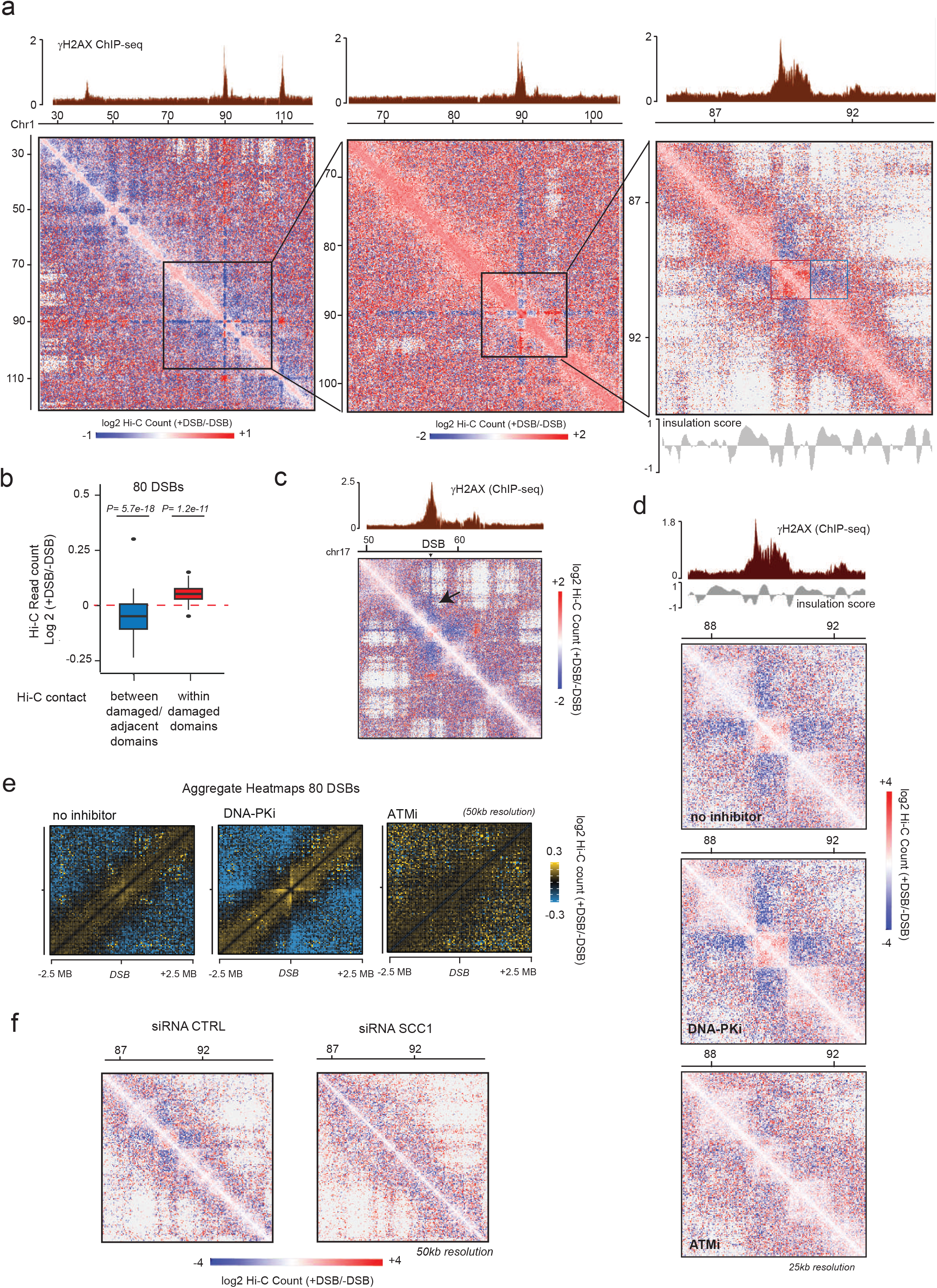
Cohesin and ATM-dependent TAD reinforcement in response to DSBs. (a) Hi-C contact matrix of the log2 (+DSB/-DSB) in DIvA cells. A region of the chromosome 1 is shown at three different resolutions: 250 kb (left panel), 100 kb (middle panel) and 25 kb (right panel). The γH2AX ChIP-seq signal following DSB induction is shown on the top panel and indicates the DSBs position. The red square highlights a damaged TAD, within which *cis* interactions are enhanced, while the blue square highlights decreased interaction between the damaged TAD and its adjacent TAD. One representative experiment is shown. (b) Boxplot showing the differential Hi-C read counts (as (log2 +DSB/-DSB)) within γH2AX domains containing the 80 best induced DSBs (red) or between these 80 damaged domains and their adjacent chromatin domains (blue). P-values, non-parametric wilcoxon test tested against μ=0. (c) Hi-C contact matrix of log2 (+DSB/-DSB) on a region located on chromosome 17 at 50 kb resolution. The contacts engaged by the DSB itself are indicated with a black arrow. γH2AX ChIP-seq track (+DSB) is shown on the top panel. One representative experiment is shown. (d) Hi-C contact matrix of the log2(+DSB/-DSB) without inhibitor (top panel), with DNA-PK inhibitor (middle panel) or with ATM inhibitor (bottom panel). A damaged region of the chromosome 1 is shown at a 25 kb resolution. Grey track represents the insulation score pre-existing to DSB induction (from Hi-C –DSB) (e) Averaged Hi-C contact matrix of the log2 (+DSB/-DSB) in untreated cells (left panel), upon DNA-PK inhibition (middle panel) or upon ATM inhibition (right panel), centered on the 80 best-induced DSBs (50 kb resolution on a 5 Mb window). (f) Hi-C contact matrix of the log2(+DSB/-DSB) on a region located on chromosome 1 at a 50 kb resolution in DIvA cells transfected with a control siRNA or a siRNA directed against SCC1.

We further investigated the contribution of PI3-Kinases involved in response to DSB by performing Hi-C in presence of inhibitors of ATM and DNA-PK, which respectively negatively and positively impact γH2AX accumulation at DSBs (in contrast to ATR inhibition, which does not noticeably alter γH2AX foci formation in DIvA cells)^5,17^. Notably, DNA-PK inhibition exacerbated the increase in intra-TAD contacts following DSB induction, while ATM inhibition abrogated it (Fig. 1d, Fig. S1a). TAD structures visualized on Hi-C maps are believed to arise thanks to cohesin-mediated loop extrusion^18^. Our previous work indicated that a bidirectional, divergent, cohesin-dependent loop-extrusion process takes place at DSBs^5^. This DSB-anchored loop extrusion can be visualized on differential Hi-C maps by a “cross” pattern centered on the DSB (Fig. 1e). Notably, ATM inhibition impaired loop extrusion, while DNA-PK inhibition strongly increased it (Fig. 1e). Moreover, depletion of the cohesin subunit SCC1, which abolishes DSB-induced loop extrusion^5^, decreased the reinforcement of intra TAD-contacts in damaged, γH2AX-decorated, chromatin domains (Fig. 1f, Fig. S1b).

Altogether these data indicate that ATM triggers cohesin-mediated loop extrusion arising from the DSB and the insulation of the damaged TADs from the surrounding chromatin.

### ATM drives clustering of damaged TADs, in a cell cycle regulated manner

We further analyzed Hi-C data with respect to long-range contacts within the nuclear space. Hi-C data revealed that DSBs cluster together (Fig. 2a, red square away from the diagonal), as previously observed using Capture Hi-C^9^. The higher resolution of this Hi-C dataset now enables us to conclude that DSB clustering takes place between entire γH2AX-decorated TADs and can happen between DSBs induced on the same chromosome (Fig. S2a) as well as on different chromosomes (Fig. S2b). Of interest, some γH2AX domains were able to interact with more than a single other γH2AX domain (Fig. 2b, black arrows). Notably, this ability to form clusters of multiples TADs (also known as TADs cliques^19^) upon DSB induction correlated with several DSB-induced chromatin features that occur at the scale of an entire TAD^3^, including γH2AX, 53BP1 and ubiquitin chains levels as well as the depletion of histone H1 around DSB detected by ChIP-seq (Fig. 2c). Moreover, it also correlated with initial RNAPII occupancy prior DSB induction indicating that DSBs prone to cluster and form damaged TAD cliques are those occurring in transcribed loci (Fig. 2c).

**Figure 2:**
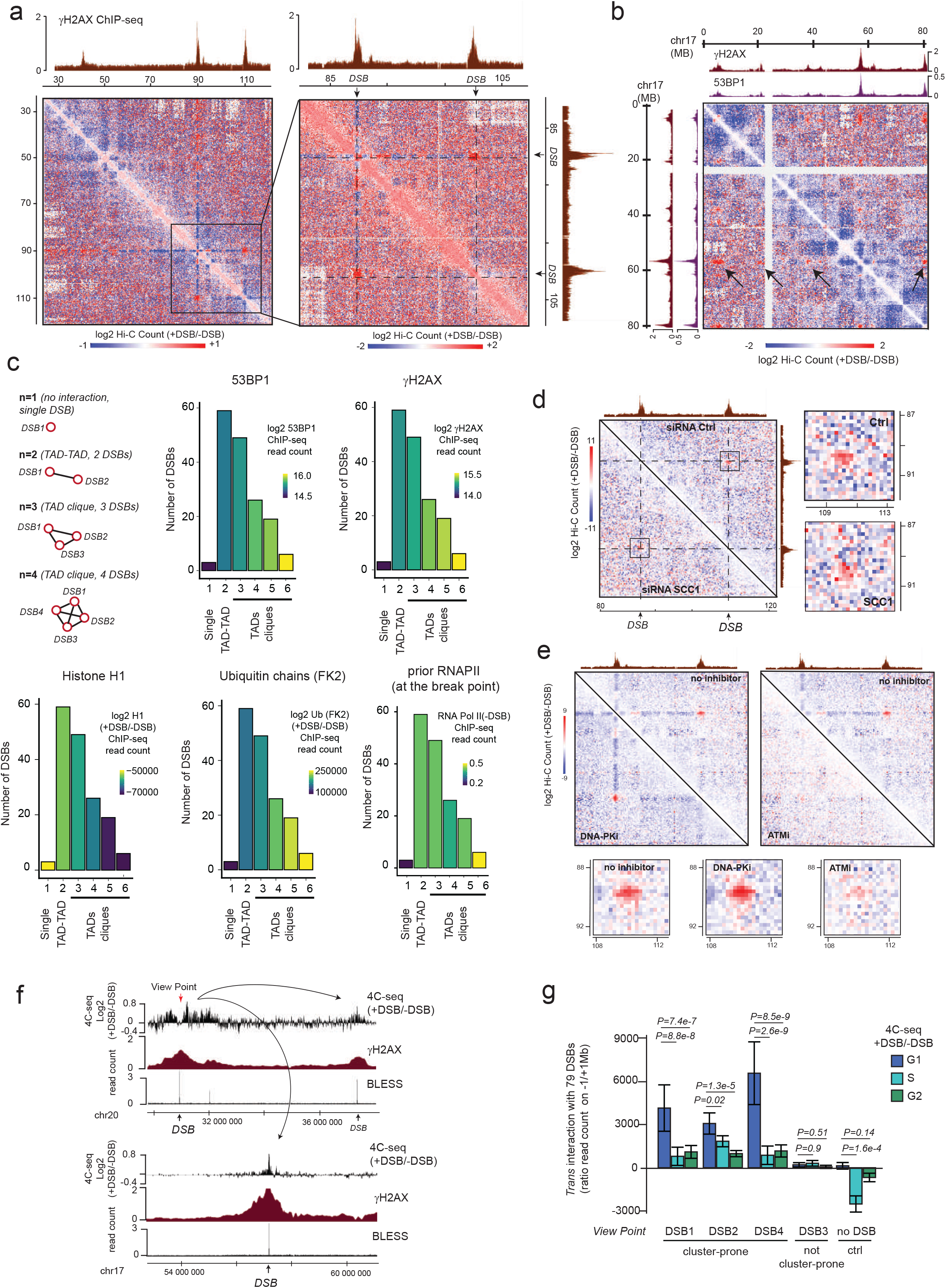
Cell cycle regulated, ATM-dependent but cohesin- and DNA-PK-independent clustering of damaged-TADs. (a) Hi-C contact matrix of the log2 (+DSB/-DSB) on a region of the chromosome 1 at two different resolutions: 250 kb (left panel) and 100 kb (right panel). γH2AX ChIP-seq track following DSB induction is shown on the top panel and on the right. One representative experiment is shown. (b) Hi-C contact matrix of the log2 (+DSB/-DSB) on a region of the chromosome 17 at 250 kb resolution. γH2AX and 53BP1 ChIP-seq tracks following DSB induction are shown on the top panel and on the left. The black arrows indicate clustering of one DSB on the chromosome 17, with several other DSBs on the same chromosome. One representative experiment is shown. (c) γH2AX domains were categorized based on their propensity to not interact with any other γH2AX domain (single), with one other γH2AX domain (TAD-TAD) or with multiple other γH2AX domains (TAD cliques containing 3 to 6 DSBs). ChIP-seq levels of γH2AX (+DSB), 53BP1 (+DSB), H1 (log2 +DSB/-DSB), Ubiquitin chains detected with the FK2 antibody (log2 +DSB/-DSB) or pre-existing RNAPII (−DSB) within the corresponding domains were computed across each category. (d) Left panel: Hi-C contact matrix of the log2(+DSB/-DSB) upon Ctrl (upper right) or SCC1 depletion (lower left). A region of the chromosome 1 is shown at 250 kb resolution. The γH2AX ChIP-seq track following DSB induction is shown on the top and on the right. Right panel: magnification of the black square, showing Hi-C contacts between the two γH2AX domains. (e) Hi-C contact matrix of the log2 (+DSB/-DSB) without inhibitor, with a DNA-PK inhibitor or with an ATM inhibitor as indicated. A region of the chromosome 1 is shown with a 250 kb resolution. γH2AX ChIP-seq track following DSB induction is shown on the top. Bottom panel: magnification, showing Hi-C contacts between the two γH2AX domains. (f) Genomics tracks showing differential 4C-seq (log2 (+DSB/-DSB)) (smoothed with a 10 kb span) obtained using a DSB located on chr20 as a viewpoint (red arrow), γH2AX ChIP-seq and BLESS, on a ∼8 Mb window of chromosome 20 (top panel) and on a ∼8 Mb window of chromosome 17 (bottom panel). Black arrows represent interactions between the DSB targeted by the viewpoint and two other DSBs, one located on the same chromosome (chr20) and one located on another chromosome (chr17). One representative experiment is shown. (g) *Trans* interactions (log2 ratio +DSB/-DSB) between the view point and the other DSBs (n=79) were computed from 4C-seq experiments in synchronized cells (G1, S and G2 as indicated). Three cluster-prone DSBs, one not cluster-prone and one control undamaged locus were used as viewpoints. *P*, non-parametric paired wilcoxon test.

We further examined the effect of cohesin depletion on damaged TAD clustering. Inspection of individual DSBs indicated that SCC1 depletion by siRNA did not alter clustering (Fig. 2d). Quantification of *trans* interactions between all DSBs also indicates that SCC1 depletion did not modify the ability of damaged TAD to physically interact together (Fig. S2c). Additionally, we found that inhibition of ATM compromised DSB clustering, whilst inhibiting DNA-PK activity triggered a substantial increase in DSB clustering (Fig. 2e, Fig. S2d).

Given the conflicting data regarding the cell cycle regulation of DSB clustering^8,9,12^, we further investigated DSB clustering in synchronized cells. DSB clustering (*i*.*e*. damaged TAD-TAD interaction) could be readily detected by 4C-seq when using a DSB as a view point, as shown by the increase of 4C-seq signal observed on other DSBs induced on the genome (Fig. 2f). We used five individual view-points: one control view point located on an undamaged locus, and four viewpoints at DSBs sites, three of which being “cluster-prone” DSBs, and one efficiently induced DSB which is unable to cluster with other DSBs. 4C-seq experiments performed before and after DSB induction in synchronized cells indicated that DSB clustering is readily detectable during G1 and is strongly reduced during the other cell cycle stages (see an example Fig. S2e). G1-specific DSB clustering was observed only when using as viewpoints “clustering-prone” DSBs, but not when using the undamaged control locus or the DSB unable to cluster (Fig. 2g).

Taken altogether, our results indicate that upon DSB formation, TADs that carry DSBs are able to physically contact each other in the nuclear space (*i*.*e*. cluster) in a manner that is entirely dependent on ATM, exacerbated upon DNA-PK inhibition, and mostly independent of the cohesin complex. Damaged TAD clustering mostly takes place in G1 and correlates with TAD-scale DSB-induced chromatin modifications (γH2AX, Ubiquitin accumulation and H1 depletion) as well as 53BP1 accumulation.

### A new “D” sub-compartment forms following DSB induction

Previous work identified the existence of two main, spatially distinct, self-segregated, chromatin “compartments” in mammalian nuclei. These chromatin compartments were determined by Principal Component Analysis (PCA) of Hi-C chromosomal contact maps where the first principal component allowed to identify loci that share similar interaction pattern, and that can be visualized linearly using eigenvectors. Further correlations with epigenomic features revealed that these two spatially segregated compartments correspond to active (the “A” compartment or euchromatin) and inactive chromatin (the “B” compartment or heterochromatin)^20^. The identification of A/B compartment using our Hi-C datasets revealed that DSB induction does not trigger major changes in genome compartmentalization into euchromatin *versus* heterochromatin (Fig. S3a). Saddle plots further confirmed that neither DSB treatment nor the pharmacological inhibition of DNAPK and ATM significantly modified the ability of the genome to segregate into active A and inactive B compartments (Fig. S3b). Moreover, DSB induction did not generally lead to compartment switch of the underlying chromatin domain, except in very few cases: Among the 80 DSBs induced by AsiSI, 58 DSBs were induced in the A compartment and all of them remained in the A compartment following DSB induction (see an example Fig. S3c top panel). Conversely, among the 22 DSBs induced in the B compartment, only 4 showed a shift from B to A (see two examples Fig. S3c middle and bottom panels). We further investigated the relationship between the compartment type and the ability of DSBs to cluster together. Of interest, DSB clustering was detectable mostly for DSBs in the A compartment (Fig. S3d).

Beyond the main classification between A/B compartments, sub-compartments have since been identified using higher resolution Hi-C maps, which correspond to subsets of heterochromatin loci (B1-B4) and of active loci (A1-A2)^21^. Of interest, such sub-compartments also correspond to microscopically visible nuclear structures such as nuclear speckles (A1)^22^ or Polycomb bodies (B1)^21^ for instance. Given that previous studies have long identified large, microscopically detectable γH2AX bodies following DNA damage and that our Hi-C data revealed clustering of damaged TADs, we postulated that DSBs may also induce a sub-compartment, in particular within the A compartment (*i*.*e*,: some A compartment, damaged-loci further segregate from the rest of the active compartment). In order to investigate this point, we applied PCA analysis on differential Hi-C maps (*i*.*e*. contact matrices of +DSB/-DSB) on each individual chromosome. The first Chromosomal Eigenvector (CEV, PC1) allowed us to identify a DSB-induced chromatin compartment mainly on chromosomes displaying a large number of DSBs (chr1,17 and X) (Fig. S4a, Fig. 3a,). Notably, a similar analysis on Hi-C maps generated upon DNA-PK inhibition, which impairs repair^17^ and increases DSB clustering (Fig. 2), allowed to identify this compartment on more chromosomes (such as chr6 for instance, Fig. S4b, bottom track). This sub-compartment displayed a very strong correlation with γH2AX-decorated chromatin following DSB (Fig. 3a, Fig. S4a-d) and was henceforth further named “D” sub-compartment (for DSB-induced compartment). Yet, further inspection revealed that the D sub-compartment is not solely generated through the clustering of damaged chromatin (*i*.*e*. TADs that carry DSBs and are enriched in γH2AX). Indeed, we could identify chromatin domains, not containing any DSB and not decorated by γH2AX, that associate with the D sub-compartment after damage (blue rectangle Fig. 3b). After exclusion of γH2AX-covered chromatin domains, correlation analysis using chromosomes 1,17 and X, on which the D sub-compartment was readily detected, indicated that non-damaged loci that tend to segregate with the D compartment are enriched in H2AZac, H3K4me3 and H3K79me2 (Fig. S4e, Fig. 3b). Conversely, these loci targeted to the D compartment displayed a negative correlation with repressive marks such as H3K9me3 (Fig. S4e). A similar trend was observed when D sub-compartment was computed from the Hi-C data obtained in presence of the DNA-PK inhibitor and correlation analysis performed on all chromosomes showing D compartmentalization (*i*.*e*, chr 1,2,6,9,13,17,18,20 and X) (Fig. S4e bottom panel). Altogether our data indicate that upon DSB production on the genome, damaged TADs, covered by γH2AX/53BP1, form a new chromatin compartment that segregates from the rest of the genome and in which some additional undamaged loci that exhibit chromatin marks typical of active transcription can be further targeted.

**Figure 3.**
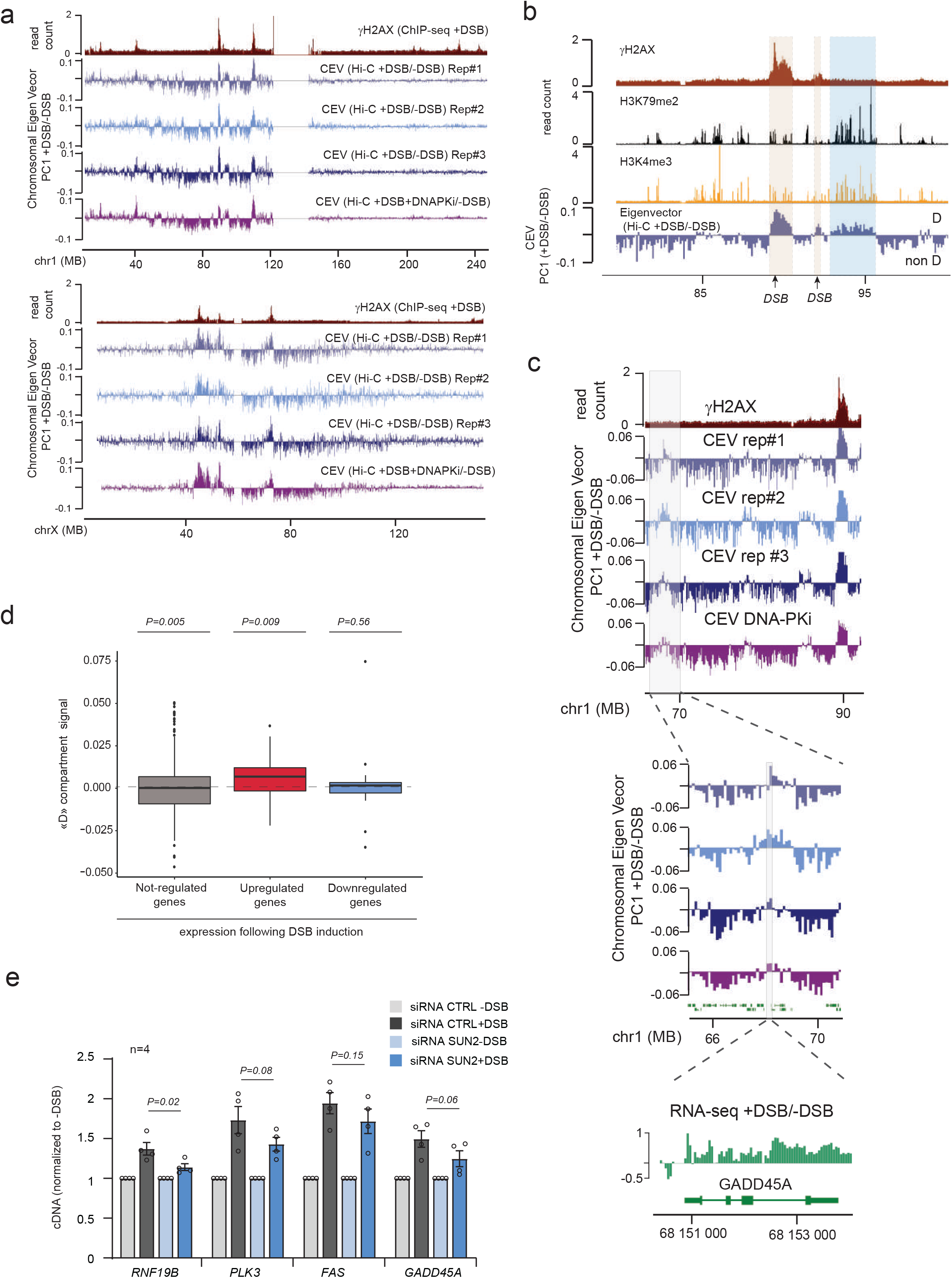
Formation of a DSB-specific sub-compartment that ensures optimal activation of the DDR. (a) Genomic tracks of γH2AX ChIP-seq and first Chromosomal eigenvector (CEV) computed on differential (+DSB/-DSB) Hi-C matrix on chromosome 1 (top panel) and chromosome X (bottom panel). Three biological replicate experiments are shown as well as the CEV obtained upon DNA-PK inhibition. (b) Genomic tracks of γH2AX (red), H3K79me2 (black) and H3K4me3 (yellow) ChIP-seq, and the first Chromosomal Eigenvector computed on the differential Hi-C (CEV, blue). The brown rectangles highlight genomic regions present in D sub-compartment that carry a DSB and are enriched in γH2AX. In contrast the blue rectangle shows a genomic region that is devoid in γH2AX and DSB, but is nevertheless found in the D sub-compartment. (c) As in (a) but with a zoom on an undamaged region of the chromosome 1 that displayed positive D sub-compartment signal. The differential RNA-seq (log2 (+DSB/-DSB)) for this region containing the p53-target gene GADD45A is also shown (green). (d) Boxplot showing the quantification of the D compartment signal computed from Hi-C data (+DSB+DNA-PKi/-DSB) on genes that are not regulated following DSB induction (Not-regulated genes, grey), genes that are upregulated following DSB induction (Upregulated genes, red) or genes that are downregulated following DSB induction (Downregulated genes, blue), identified by RNA-seq. (e) RT-qPCR quantification of the expression level of four genes (*RNF19B, FAS, PLK3* and *GADD45A*) before and after DSB induction in cells transfected with control or SUN2 siRNA. n=4 independent experiments.

### A subset of DNA damage responsive genes segregates with the D sub-compartment to achieve optimal activation

In order to decipher the nature of the active genes targeted to the D compartment, we further explored the DNA motifs enriched on “D” genes compared to “non D” genes, *i*.*e*. genes recruited to the D compartment, *versus* the one that do not display targeting to the D compartment (discarding all genes directly comprised in γH2AX domains). Notably, the top enriched motifs included OSR1, TP73, Nkx3.1 and E2F binding sites, which are tumor suppressor and /or known to be involved in the DNA damage response (Fig. S4f)^23–26^, suggesting a direct physical targeting of DNA damage responsive genes to the “D” sub-compartment. In agreement, visual inspection revealed that some known p53 target genes which are upregulated following DSB induction were associated with the D compartment, even when as far as >20MB from the closest DSB (see an example Fig. 3c). To test the hypothesis that DNA damage responsive genes are recruited to the D compartment, we performed RNA-seq before and after DSB induction and retrieved genes that are upregulated following DSB induction. Notably, genes upregulated following DSB induction displayed a higher D compartment signal compared to genes that were either not regulated or downregulated after DSBs (Fig. 3d). Of note, if some of the upregulated genes were indeed targeted to the D compartment, this was not the case for all of them. Importantly, the upregulated genes targeted to the D-compartment were not in average closer to DSBs than the upregulated genes not-targeted to the D compartment (Fig. S4g), ruling out a potential bias due to the genomic distribution of AsiSI DSBs.

In order to determine whether recruitment of those genes to the D sub-compartment contribute to their activation following DNA damage, we investigated the consequence of disrupting DSB clustering (and hence formation of D compartment) by depleting the SUN2 component of the LINC complex, previously found as a DSB-clustering promoting factor^9,11^. SUN2 depletion altered the transcriptional activation of genes found to be upregulated and targeted to the D sub-compartment upon DSB in DIvA cells (Fig. 3e).

Altogether these data indicate that DSB induction triggers the formation of a novel chromatin sub-compartment that comprises not only damaged TADs, decorated by γH2AX and 53BP1, but also a subset of genes upregulated following DNA damage, for which targeting to D sub-compartment is required for optimal activation. Altogether this suggests a role of the D sub-compartment, and hence DSB clustering, in the activation of the DNA Damage Response.

### DSB-induced reorganization of chromosome folding favors translocations

Importantly, while our above data suggest a beneficial role of DSB clustering in potentiating the DDR, it may also be detrimental, since bringing two DSBs in a close proximity may fosters translocations (illegitimate rejoining of two DSBs), as previously proposed^10^. We therefore assessed by qPCR the frequency of translocations events occurring in DIvA cells post-DSB induction, in conditions where we found altered DSBs clustering and D compartment formation.

Notably, translocations are increased in G1 compared to S/G2-synchronized cells (Fig. 4a), in agreement with an enhanced DSB clustering observed in G1 cells (Fig. 2). Moreover, DNA-PK inhibition, that increased D-compartment formation (Fig. 2e, Fig. S2d, Fig. S4b) also strongly increased translocation frequency (Fig. 4b). On another hand, depletion of 53BP1 (Fig. S5a), previously found to mediate repair foci phase separation^13^, as well as a treatment with 1,6-hexanediol, which disrupts phase condensates (Fig. S5b), decreased translocations (Fig. 4c). Similarly, depletion of SUN2, member of the LINC complex and of ARP2, an actin branching factor (Fig. S5a), reported as mediating DSB clustering^9,11,12^, decreased translocations (Fig. 4c). Surprisingly, depletion of the cohesin subunits SMC1 or SCC1 also decreased translocation frequency (Fig. 4d, Fig S5c). This was unexpected since SCC1-depleted cells do not display clustering defects (Fig. 2).

**Figure 4.**
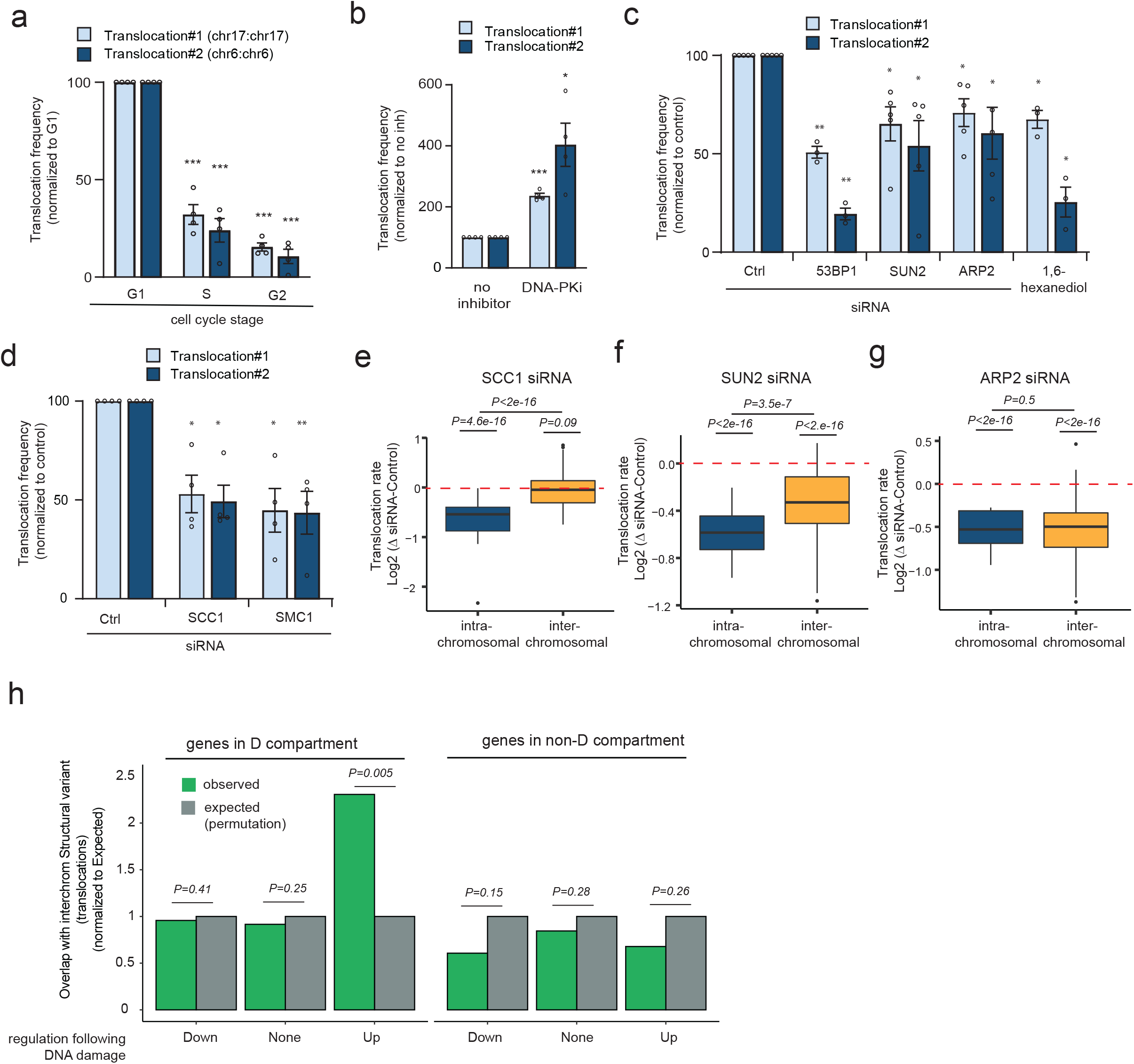
DSB-induced loop extrusion and D-compartment formation drive translocations. (a) qPCR quantification of translocations frequency for two independent translocations following DSB induction in cells synchronized in the G1, S or G2 phase (n=4 independent replicates). *P*= paired t-test, * *P*<0.05, ** *P*<0.001, \*\*\**P*<0.0005 (b) qPCR quantification of translocations frequency for two independent translocations following DSB induction with or without DNA-PK inhibitor (n=4 independent replicates). (c) qPCR quantification of translocations frequency for two independent translocations following DSB induction in Control, 53BP1, SUN2 or ARP2 depleted cells or upon 1,6-Hexanediol treatment (n≥3 independent replicates). (d) As in (c) but upon Control, SMC1 or SCC1 depletion (n=4 independent replicates). (e) Intra-chromosomal (blue) or inter-chromosomal translocations (yellow) were quantified using multiplexed amplification followed by high throughput sequencing (amplicon-seq) between 20 different DSBs induced in DIvA cell line, upon Ctrl or SCC1 depletion (log2 siSCC1/siCTRL) (n=4 independent replicates). P-values, non-parametric wilcoxon test tested against μ=0. intra vs inter-chromosomal, *P=*paired wilcoxon test. (f) As in (e) but the quantification was performed in SUN2 depleted cells (n=4 independent replicates). (g) As in (e) but the quantification was performed in ARP2 depleted cells (n=4 independent replicates). (h) Observed (green) and expected (obtained through 1000 permutations) overlap between breakpoint positions of inter-chromosomal translocations identified on cancer genomes and genes targeted to the D compartment, either upregulated, downregulated or not regulated following DSB induction (identified by RNA-seq) as indicated, compared to their counterparts not targeted to the D compartment.

Given that the two translocations assessed by our qPCR assay are both intra-chromosomal translocations (*i*.*e*.: rejoining of two distant DSBs located on the same chromosome) we hypothesized that translocation frequency at the intra-chromosomal level may also be regulated by the DSB-induced loop extrusion that depends on the cohesin complex. In order to investigate more broadly translocation events between multiple DSBs induced in the DIvA cell line, we designed a novel multiplexed amplification protocol followed by NGS sequencing. In control cells, we could readily detect increased translocation frequency upon induction of DSB compared to control genomic locations (Fig. S5d). Strikingly, depletion of SCC1 decreased the frequency of intra-chromosomal translocations, while leaving inter-chromosomal translocations unaffected (Fig. 4e). In contrast depletion of SUN2 and ARP2 decreased both intra- and inter-chromosomal translocations (Fig. 4f-g). Taken together these data suggest that both the DSB-induced loop extrusion and the formation of the D sub-compartment through clustering of damaged TADs, display the potential to generate translocations.

Given our above finding that a subset of genes upregulated following DSB induction can be physically targeted to the D compartment after break induction (Fig. 3), we further hypothesized that such a physical proximity may account for some of the translocations observed on cancer genomes. We retrieved breakpoint positions of inter-chromosomal translocations of 1493 individuals across 18 different cancers types (from^27^), and assessed their potential overlap with genes targeted to the D sub-compartment (reproducibly detected in the three Hi-C replicates on chr1,17 and X, on which D sub-compartment could be identified accurately). D-targeted genes were further sorted as either upregulated, downregulated or not significantly altered following

DSB induction, and compared to their counterparts not targeted to the D compartment. We found that genes that are upregulated following DSB induction and that are targeted to the D compartment displayed a significant overlap with translocations breakpoints, in contrast to genes that are not targeted to the D compartment (non-D) (Fig. 4h). Altogether these data indicate that the relocalization of upregulated genes during the DNA Damage response in the DSB-induced sub-compartment likely accounts for some of the translocations detected on cancer genomes. Given that DDR genes comprise a number of tumor suppressor genes, such a physical proximity of these genes with DSBs within the D sub-compartment formed in response to DNA damage, may be a key mechanism driving oncogenesis, through fostering the instability of tumor suppressor genes.

### Conclusion

Altogether this work shows that DSB-induced changes in chromosome architecture is an integral component of the DNA Damage Response, but also acts as a double-edged sword that can challenge genomic integrity through the formation of translocations.

Our data suggest that a chromatin sub-compartment arises when γH2AX/53BP1-decorated domains, established by ATM-induced loop extrusion post DSB, self-segregate from the rest of chromatin. This may, at least in part, occur thanks to the LLPS properties of 53BP1^13,14,28^. This DSB-induced (“D”) sub-compartment further recruits a subset of genes involved in the DNA damage response and contributes to their activation (Fig. S5e). This model is in agreement with previous work which identified 53BP1 as critical for p53 target genes activation^29^, with the findings that disrupting 53BP1 droplet formation alters checkpoint activation^13^ and with the fact that enhanced 53BP1 phase separation triggers an elevated p53 response^30^ as does the loss of TIRR, a protein that regulates 53BP1 association to DSBs^31,32^. We propose that the formation of the “D” sub-compartment allows to precisely tune the magnitude of the DDR with respect to DSB load and persistency, providing a function for these enigmatically large γH2AX/53BP1-decorated chromatin domains and to DSB clustering. Furthermore, this observation may provide a rationale for why so many transcription factors (including p53) were found recruited at DSBs repair foci^33^. While initially thought to allow chromatin remodeling in order to enhance DSB repair, the recruitment of transcription factors to DSB repair foci may in fact rather reflects the relocalization of DDR genes within the D compartment (hence at physical proximity of the DSB).

Yet, this comes at the expense of potential translocations, as both loop extrusion and coalescence of damaged TAD are able to bring linearly distant DSBs in close physical proximity (Fig. S5e). Importantly, we found that the genes upregulated in response to DSB and relocated to the D compartment displayed significant overlap with translocation breakpoints identified by whole genome sequencing in patient cancer samples. In agreement with an increased occurrence of structural variants on tumor suppressor genes^27^, we propose that the physical targeting of DNA damage responsive genes to the D compartment, by bringing DSBs and DDR genes in close spatial proximity, may occasionally trigger deleterious rearrangements on genes involved in the control of cell proliferation and apoptosis upon DNA damage, and may hence act as a critical driver of oncogenesis by disrupting the integrity of tumor suppressor genes.

## Methods

### Cell culture and treatments

DIvA (AsiSI-ER-U20S)^16^ and AID-DIvA (AID-AsiSI-ER-U20S)^34^ cells were grown in Dubelcco’s modified Eagle’s medium (DMEM) supplemented with 10% SVF (Invitrogen), antibiotics and either 1 µg/mL puromycin (DIvA cells) or 800 µg/mL G418 (AID-DIvA cells) at 37 °C under a humidified atmosphere with 5% CO2. To induce DSBs, cells were treated with 300nM 4OHT (Sigma, H7904) for 4 h. For ATM or DNA-PK inhibition, cells were pretreated for 1 h respectively with 20μM KU-55933 (Sigma, SML1109) or 2μM NU-7441 (Selleckchem, S2638) and during subsequent 4OHT treatment. Treatment with 10% 1,6-hexanediol (Sigma, 240117) was performed for 3 min before the end of the 4OHT treatment. For cell synchronization, cells were incubated for 18 h with 2 mM thymidine (Sigma, T1895), then released during 11 h, followed by a second thymidine treatment for 18 hr. S, G2 and G1 cells were then respectively treated with OHT at, 0, 6 or 11 h following thymidine release and harvested 4 h later. siRNA transfections were performed using the 4D-Nucleofector and the SE cell line 4D-Nucleofector X kit L (Lonza) according to the manufacturer’s instructions, and subsequent treatment(s) were performed 48 h later. siRNA transfections were performed using a control siRNA (siCTRL): CAUGUCAUGUGUCACAUCU; or using a siRNA targeting *SCC1* (siSCC1): GGUGAAAAUGGCAUUACGG; or SMC1 (siSMC1): UAGGCUUCCUGGAGGUCACAUUUAA; or 53BP1 (si53BP1): GAACGAGGAGACGGUAAUA; or SUN2 (siSUN2): CGAGCCTATTCAGACGTTTCA; or ARP2 (siARP2): GGCACCGGGUUUGUGAAGU.

### Translocation assay

Translocation assays after siRNA transfection or 1,6-Hexanediol treatment were performed at least in triplicates in AID-DIvA cells as described in^35^. Translocation assay in synchronized cells was performed in DIvA cells following a 4OHT treatment (n=4 biological replicates). Two different possible translocations between different AsiSI sites were assessed by qPCR using the following primers: Translocation1_Fw: GACTGGCATAAGCGTCTTCG, Translocation1_Rev: TCTGAAGTCTGCGCTTTCCA, Translocation2_ Fw: GGAAGCCGCCCAGAATAAGA, Translocation2_Rev: TCTGAAGTCTGCGCTTTCCA. Results were normalized using two control regions, both far from any AsiSI sites and γH2AX domain using the following primers: Ctrl_chr1_82844750_Fw: AGCACATGGGATTTTGCAGG, Ctrl_chr1_82844992_Rev: TTCCCTCCTTTGTGTCACCA, Ctrl_chr17_9784962_Fw: ACAGTGGGAGACAGAAGAGC, Ctrl_chr17_9785135_Rev: CTCCATCATCGCACCCTTTG. Normalized translocation frequencies were calculated using the Bio-Rad CFX Manager 3.1 software69.

### Amplicon –seq

AID-DIvA cells were treated with or without 300nM 4OHT for 4 h followed by treatment with indole-3-acetic acid for 14 h. Cells were then lysed in cytoplasmic lysis buffer (50mM HEPES pH7.9, 10mM KCl2, 1.5mM MgCl2, 0.34M sucrose, 0.5% triton X-100, 10% glycerol, 1mM DTT) for 10 minutes on ice, then washed once in cytoplasmic lysis buffer before lysis in genomic extraction buffer (50mM Tris pH8.0, 5mM EDTA, 1% SDS, 0.5mg/mL proteinase K). Lysate was incubated at 60°C for 1 h. Genomic DNA was then ethanol precipitated on ice for 1h, pelleted at 19,000g for 20 min and washed twice in 75% ethanol. Genomic DNA was then used in a multiplex PCR reaction that amplified 25 target sites; 20 AsiSI cut sites and 5 uncut control sites (Supplementary Table 1). Amplicons were size selected using SPRIselect beads (Beckman, B23318) and subjected to DNA library preparation via the NEBNext Ultra II kit (NEB, E7645L). Libraries were pooled at equimolar concentrations and sequenced via an Illumina NextSeq 500 system using paired end 150 cycles. The data was analyzed via our custom tool mProfile, available at github.com/aldob/mProfile. This identified the genomic primers used in the original genomic PCR reaction to amplify each read in the pair. Translocated reads were therefore identified as those where each read in a pair was amplified by a different primer set, and this was normalized to the total reads that were correctly amplified by these primer sets.

### RT-qPCR

RNA was extracted from fresh DIvA cells before and after DSB induction using the RNeasy kit (Qiagen). RNA was then reverse transcribed to cDNA using the AMV reverse transcriptase (Promega, M510F). qPCR experiments were performed to assess the levels of cDNA using primers targeting RPLP0 (FW: GGCGACCTGGAAGTCCAACT; REV: CCATCAGCACCACAGCCTTC), RNF19B (FW: CATCAAGCCATGCCCACGAT; REV: GAATGTACAGCCAGAGGGGC), PLK3 (FW: GCCTGCCGCCGGTTT; REV: GTCTGACGTCGGTAGCCCG), FAS (FW: ATGCACACTCACCAGCAACA; REV: AAGAAGACAAAGCCACCCCA) or GADD45A (FW: ACGATCACTGTCGGGGTGTA; REV: CCACATCTCTGTCGTCGTCC). cDNA levels were then normalized with RPLP0 cDNA level, then expressed at the percentage of the undamaged condition.

### Immunofluorescence

DIvA cells were grown on glass coverslips and fixed with 4% paraformaldehyde during 15 min at room temperature. Permeabilization step was performed by treating cells with 0,5% Triton X-100 in PBS for 10 min then cells were blocked with PBS-BSA 3% for 30min. Primary antibodies targeting RNA PolI (Santa Cruz sc48385) or PML (Santa Cruz sc-966 (PG-M3)) were diluted 1:500 in PBS-BSA 3% and incubated with cells overnight at 4°C. After washes in 1X PBS, cells were incubated with anti-mouse secondary antibody (conjugated to Alexa 594 or Alexa 488, Invitrogen), diluted 1:1000 in PBS-BSA 3%, for 1h at room temperature. After a DAPI staining, Citifluor (Citifluor, AF-1) was used for coverslips mounting. Images were acquired with the software MetaMorph, using the 100X objective of a wide-field microscope (Leica, DM6000), equipped with a camera (DR-328G-C01-SIL-505, ANDOR Technology).

### Western Blot

Western Blot experiments were performed as in^5^ using primary antibody targeting SUN2 (Abcam ab124916 1:1000), ARP2 (Abcam ab128934 1:1000), 53BP1 (Novus Biologicals NB100-305 1:1000), SCC1 (Abcam ab992 1:500) or SMC1 (Abcam ab75819 1:1000).

### RNA-seq

RNA-seq was performed as described in^35^. RNA-seq were mapped in paired-end to a custom human genome (hg19 merged with ERCC92) using STAR. Count matrices were extracted using htseq-count with union as resolution-mode and reverse strand mode. Differential expression analysis was made on the count matrix using edgeR with two replicates per condition and differential genes were determined with log-ratio test (LRT). Whole genome coverage was computed using deeptools and bamCoverage to generate bigwig using bam files (without PCR duplicate suppression). Using a cutoff of 0.1 for the adjusted p-value and 0.5 log2 fold-change (∼41% increase/decrease of expression), we were able to determine 286 up-regulated and 125 down-regulated genes with 11 of them directly damaged by a DSB. Differential coverage between two conditions was performed using BamCompare from deeptools with setting binsize parameter at 50bp. Log2FC was calculated by edgeR in differential expression analysis.

### 4C-seq

4C-seq experiments performed in synchronized cells, before and after DSB induction were performed as in^5^. Briefly, 10-15×10^6^ DIvA cells per condition were cross-linked, lysed and digested with MboI (New England Biolabs). DNA ligation was performed using the T4 DNA ligase (HC) (Promega), and ligated DNA was digested again using NlaIII (New England Biolabs). Digested DNA was religated with the T4 DNA ligase (HC) (Promega) before to proceed to 4C–seq library preparation. 16 individual PCR reactions were performed in order to amplify ∼800ng of 4C-seq template, using inverse primers including the Illumina adaptor sequences and a unique index for each condition (Supplementary Table 2). Libraries were pooled and sent to a Nextseq500 platform at the I2BC Next Generation Sequencing Core Facility (Gif-sur-Yvette).

4C-seq data were processed as described in^5^. Briefly, bwa mem was used for mapping and samtools for sorting and indexing. A custom R script (https://github.com/bbcf/bbcfutils/blob/master/R/smoothData.R) was used to build the coverage file in bedGraph format, to normalize using the average coverage and to exclude the nearest region from each viewpoint. Differential 4C-seq data were computed using BamCompare from deeptools with binsize=50bp. Average of total Trans interactions between viewpoints and DSB were then computed using a 1Mb window around the breaks (80 best) and after exclusion of viewpoint-viewpoint (Cis) interactions.

### Hi-C

Hi-C data obtained before and after DSB induction and upon CTRL or SCC1 depletion in DIvA cells were retrieved from^5^. Hi-C experiments with or without DSB induction and upon ATM or DNA-PK inhibition were performed in DIvA cells as in^5^. Briefly, 1 million cells were used per condition. Hi-C libraries were generated using the Arima Hi-C kit (Arima Genomics) by following the manufacturer instructions. DNA was sheared to an average fragment size of 350-400 pb using the Covaris S220 and sequencing libraries were prepared on beads using the NEB Next Ultra II DNA Library Prep Kit for Illumina and NEBNext Multiplex Oligos for Illumina (New England Biolabs) following instructions from the Arima Hi-C kit.

### Hi-C data analyses

#### Hi-C heatmaps

Hi-C reads were mapped to hg19 and processed with Juicer using default settings (https://github.com/aidenlab/juicer). Hi-C count matrices were generated using Juicer at multiple resolutions: 100 kb, 50 kb, 25 kb, 10 kb and 5 kb. Hi-C heatmaps screenshots were generated using Juicebox (https://github.com/aidenlab/Juicebox/wiki/Download). Aggregate heatmaps were computed on a set of sub-matrices extracted from originals observed Hi-C matrices at 50kb resolution or 100kb resolution. Region of 5Mb around DSBs (80 best) were extracted and then averaged. Log2 ratio was then computed using Hi-C counts (+DSB/-DSB) and plotted as heatmaps.

#### Cis Contacts Quantification

For *cis* contact quantification interaction within γH2AX domains (−0.5/+0.5Mb around 80 best DSBs) were extracted from the observed Hi-C matrix at 100kb resolution, and log2 ratio was computed on damaged vs undamaged Hi-C counts (+DSB/-DSB). Adjacent windows (−1.5Mb-0.5Mb and +0.5Mb-1.5Mb around 80 best DSBs) were retrieved to quantify interactions between damaged domains and adjacent undamaged domains. Boxplots: Centre line, median; box limits, first and third quartiles; whiskers, maximum and minimum without outliers; points, outliers. Significance was calculated using non-parametric Wilcoxon test.

#### Trans contact quantification

To determine interaction changes in *trans* (inter-chromosomal) we built the whole-genome Hi-C matrix for each experiment by merging together all chr-chr interaction matrices using Juicer and R. The result is a genome matrix with 33kx33k bin interactions for 100kb resolution. Interactions between bins inside damaged TADs (240×240 for 80 DSBs) were extracted and counted for each condition, log2 ratio was calculated on normalized count (cpm), and plotted as boxplots. Boxplots: Centre line, median; box limits, first and third quartiles; whiskers, maximum and minimum without outliers; points, outliers.

#### TAD Cliques

TAD Cliques were computed using the igraph R package on an undirected graph representing DSB clustering. This graph was computed on the differential Hi-C matrix (+DSB/-DSB) counts, at 500 kb resolution, considering a change of ∼86% of interaction (0.9 in log2) as between two DSBs as a node on the graph. Averaged signal of ChIP-seq values (53BP1/ γH2AX/H1/Ubiquitin FK2) were then computed for each categories of cliques using 500kb windows around DSB. For prior RNAPII occupancy, the signal was computed on 10kb around DSBs.

#### A/B compartment

To identify the two mains chromosomal compartments (A/B), the extraction of the first eigenvector of the correlation matrix (PC1) was done on the Observed/Expected matrix at 500kb resolution using juicer eigenvector command. The resulting values were then correlated with ATAC-seq signal in order to attributes positives and negatives values to the A and B compartment, respectively, on each chromosomes. The Observed/Expected bins were arranged based on the PC1 values and aggregated into 21 percentiles, to visualize A-B interactions on our experiments (saddle plots).

#### D compartment

To identify the D compartment, we retrieved the first component (PC1) of a PCA made on the differential observed Hi-C matrix 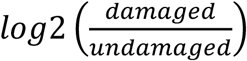 at 100kb resolution.

Each matrix was extracted from the .hic files using Juicer and the ratio was computed bin per bin. Pearson Correlation matrices were then computed for each chromosome, and PCA was applied on each matrix. The first component of each PCA was then extracted and correlated with the positions of DSB. A PC1 showing a positive correlation with DSB was then called D compartment, and PC1 showing negative correlation with DSBs were multiplied by -1. We were able to extract the D compartment on chromosomes 1,17 and X for +DSB/-DSB and chromosomes 1,2,6,9,13,17,18,20 and X for +DSB/-DSB in DNA-PKi condition. D compartment (first component of the PCA) was converted into a coverage file using rtracklayer R package. Using the same package, D compartment value was computed around DSBs and genes at 100kb resolution, and plotted as boxplot. Boxplots: Centre line, median; box limits, first and third quartiles; whiskers, maximum and minimum without outliers; points, outliers.

#### Transcription factor motif analysis

TF-binding motifs were extracted on the promoter regions (−500bp/TSS) of genes with positive value of D compartment (2161) *vs* genes with negative value (2112) using motifmatchr and TFBSTools R packages on JASPAR2020 database. Motifs were sorted by significance using fisher exact test and adjusted with Benjamini-Hochberg procedure between motifs found on gene inside the D compartment versus genes outside D compartment.

#### Translocation breakpoints

For translocation breakpoints, data from^27^ were retrieved, and only breakpoints for interchromosomal structural variant selected (N=28051). Genes reproducibly enriched in Compartment D in the three biological replicates, on chr1, 17 and X (N=604) as well as genes not enriched in Compartment D (N=1439) were retrieved. The significance of the overlap between genes and breakpoints was determined using the regioneR package^36^ using resampling test with PermTest. Briefly, we selected 1000 times a control set of genes, with same size and on the same chromosome as our original gene set. We tested the overlap between each genes and breakpoints, to determine a distribution of the number of overlaps between control set and breakpoints. We further tested if the overlap between our gene set (D compartment or non D compartment) and breakpoints was significant, by counting the number of times we got more overlap in control than in our gene set.

## Acknowledgments

We thank the genomics core facility of EMBL and of the I2BC (Centre de Recherche de Gif) for high-throughput sequencing. M.B. was supported by the CRUK Beatson Institute core grant A29252; A.B was supported by national productivity award from the MRC, MC_ST_U17040. Funding in GL laboratory was provided by grants from the European Research Council (ERC-2014-CoG 647344), Agence Nationale pour la Recherche (ANR-18-CE12-0015) and the Ligue Nationale contre le Cancer (LNCC). C.A was a recipient of a FRM fellowship (FRM FDT201904007941). T.C and N.P are INSERM researchers.

## Authors contributions

C.A., E.L., T.C., and N.P. performed and analyzed experiments. V.R., and R.M, performed bioinformatic analyses of all high-throughput sequencing datasets. A.B performed the Amplicon-seq experiment under the supervision of M.B. D.N. helped to realize and analyze 4C-seq experiments. G.L. and T.C. wrote the manuscript. All authors commented and edited the manuscript.

## Data Availability

All high-throughput sequencing data (Hi-C, 4C–seq, Amplicon-seq and RNA-seq) have been deposited to Array Express (https://www.ebi.ac.uk/arrayexpress/) under accession number E-MTAB-XXXX.

## Code availability

Source codes are available from https://github.com/LegubeDNAREPAIR/

