## Supplementary figures and legends for "ATM-dependent formation of a novel chromatin compartment regulates the Response to DNA Double Strand Breaks and the biogenesis of translocations"

Figure S1-S5

Supplemental Figures legends

Supplementary Table 1-2

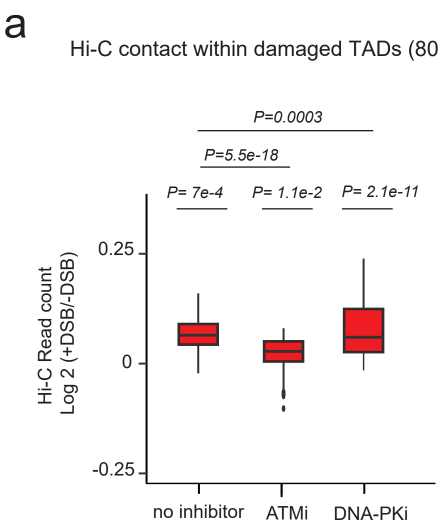
a
Hi-C contact within damaged TADs (80 DSBs)
P=0.0003
P=5.5e-18
P= 7e-4
P= 1.1e-2
P= 2.1e-11
Hi-C Read count
Log 2 (+DSB/-DSB)
0.25
0
-0.25
no inhibitor
ATMi
DNA-PKi

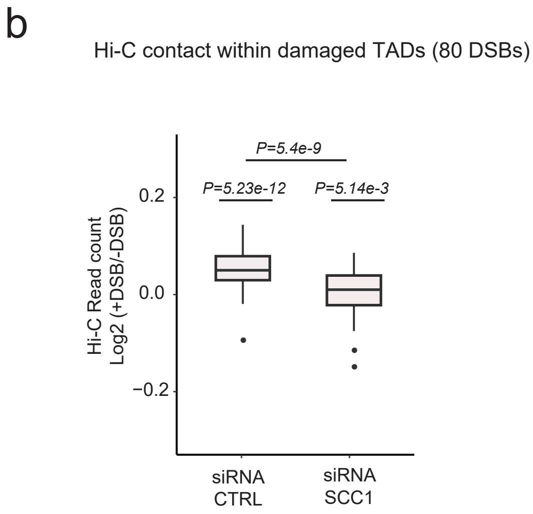
b
Hi-C contact within damaged TADs (80 DSBs)
P=5.4e-9
P=5.23e-12
P=5.14e-3
Hi-C Read count
Log2 (+DSB/-DSB)
0.2
0.0
−0.2
siRNA
CTRL
siRNA
SCC1

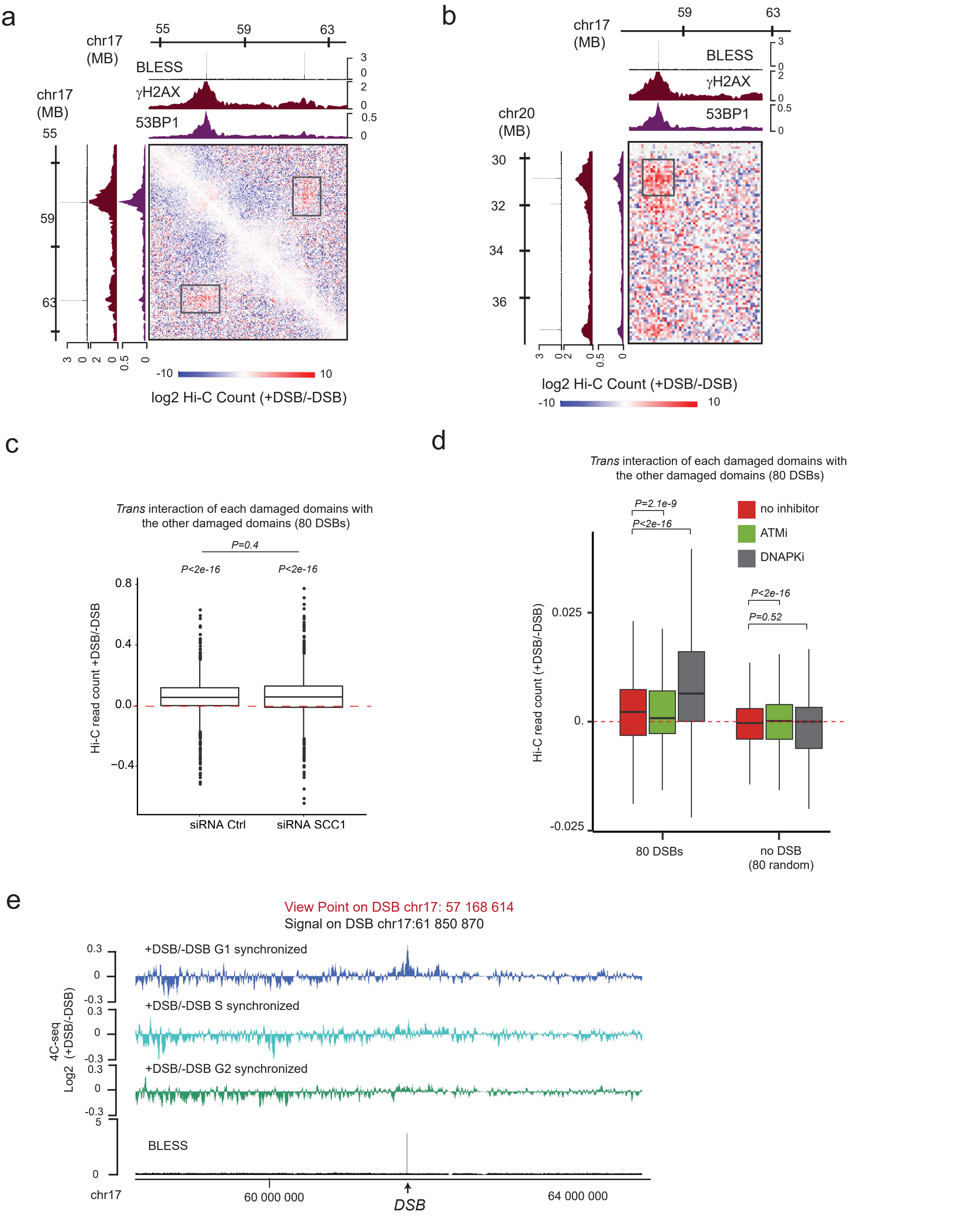
a
chr17 (MB)
55
59
63
BLESS
γH2AX
53BP1
chr17 (MB)
55
59
63
-10
10
log2 Hi-C Count (+DSB/-DSB)
b
chr17 (MB)
59
63
BLESS
γH2AX
53BP1
chr20 (MB)
30
32
34
36
log2 Hi-C Count (+DSB/-DSB)
-10
10
c
Trans interaction of each damaged domains with the other damaged domains (80 DSBs)
P=0.4
P<2e-16
P<2e-16
Hi-C read count +DSB/-DSB
siRNA Ctrl
siRNA SCC1
d
Trans interaction of each damaged domains with the other damaged domains (80 DSBs)
P=2.1e-9
P<2e-16
no inhibitor
ATMi
DNAPKi
P<2e-16
P=0.52
Hi-C read count (+DSB/-DSB)
80 DSBs
no DSB (80 random)
e
View Point on DSB chr17: 57 168 614
Signal on DSB chr17:61 850 870
+DSB/-DSB G1 synchronized
+DSB/-DSB S synchronized
+DSB/-DSB G2 synchronized
4C-seq Log2 (+DSB/-DSB)
BLESS
chr17
60 000 000
64 000 000
DSB

Arnould, Rocher et al, Figure S3

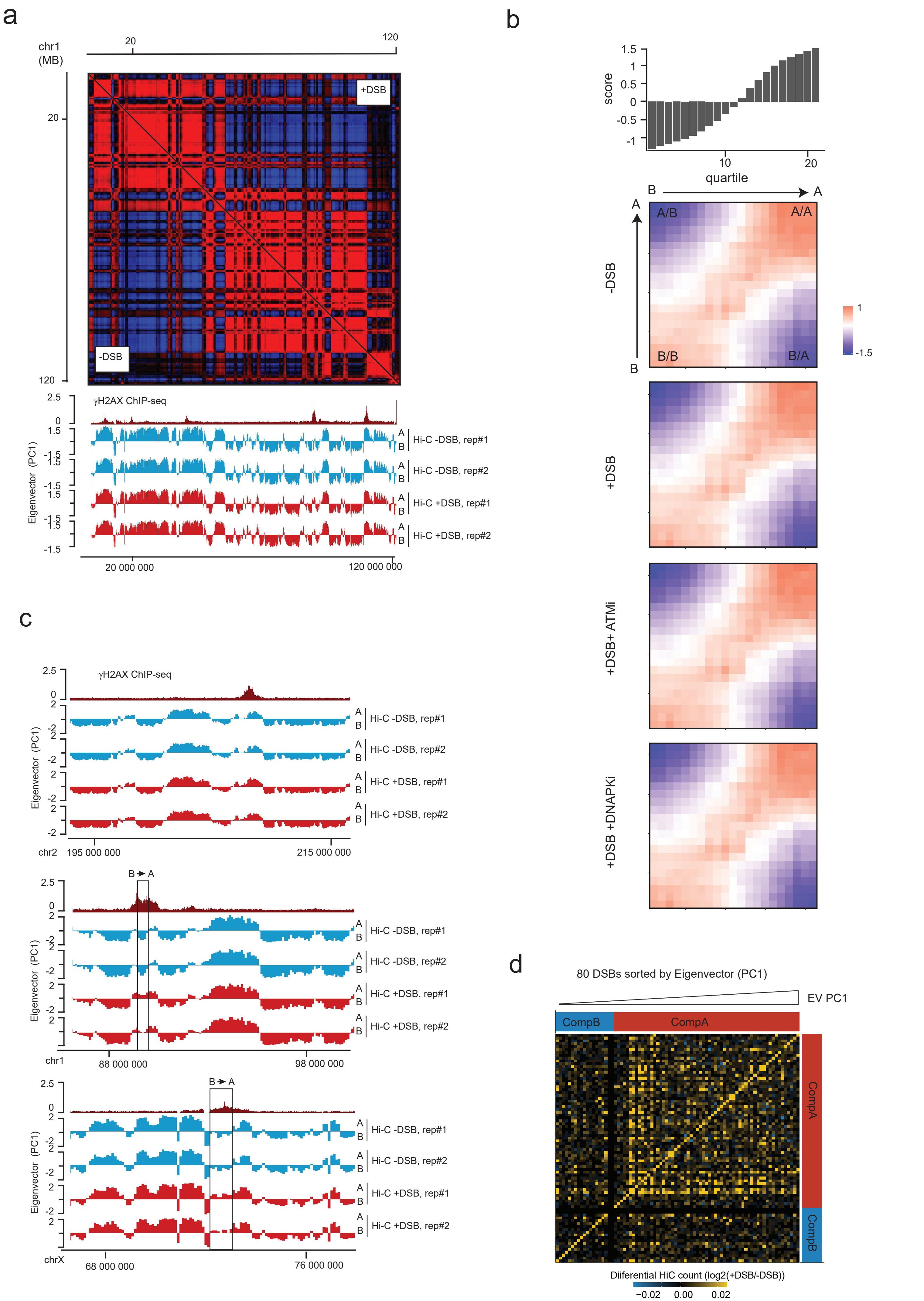

### Arnould, Rocher et al, Figure S4

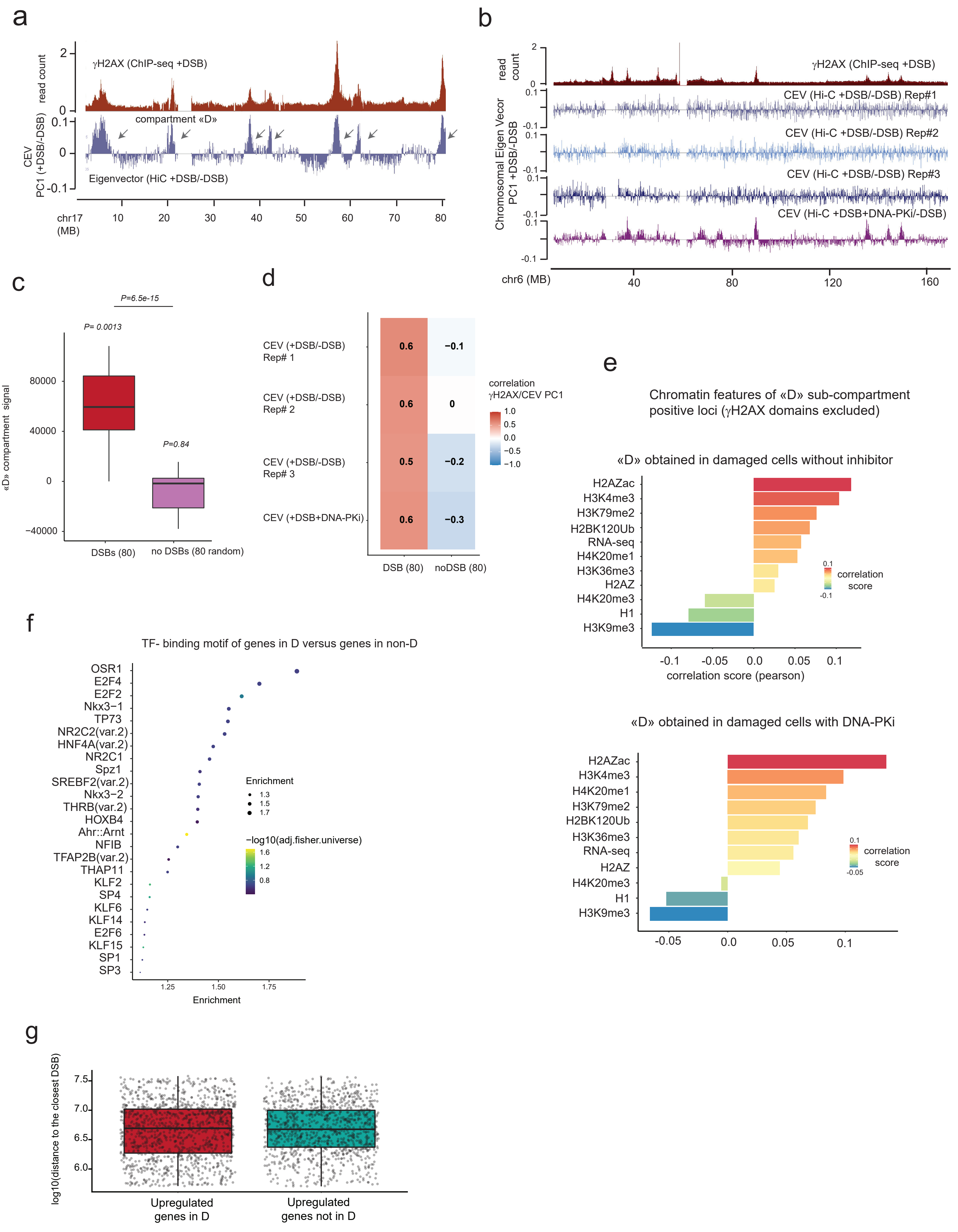

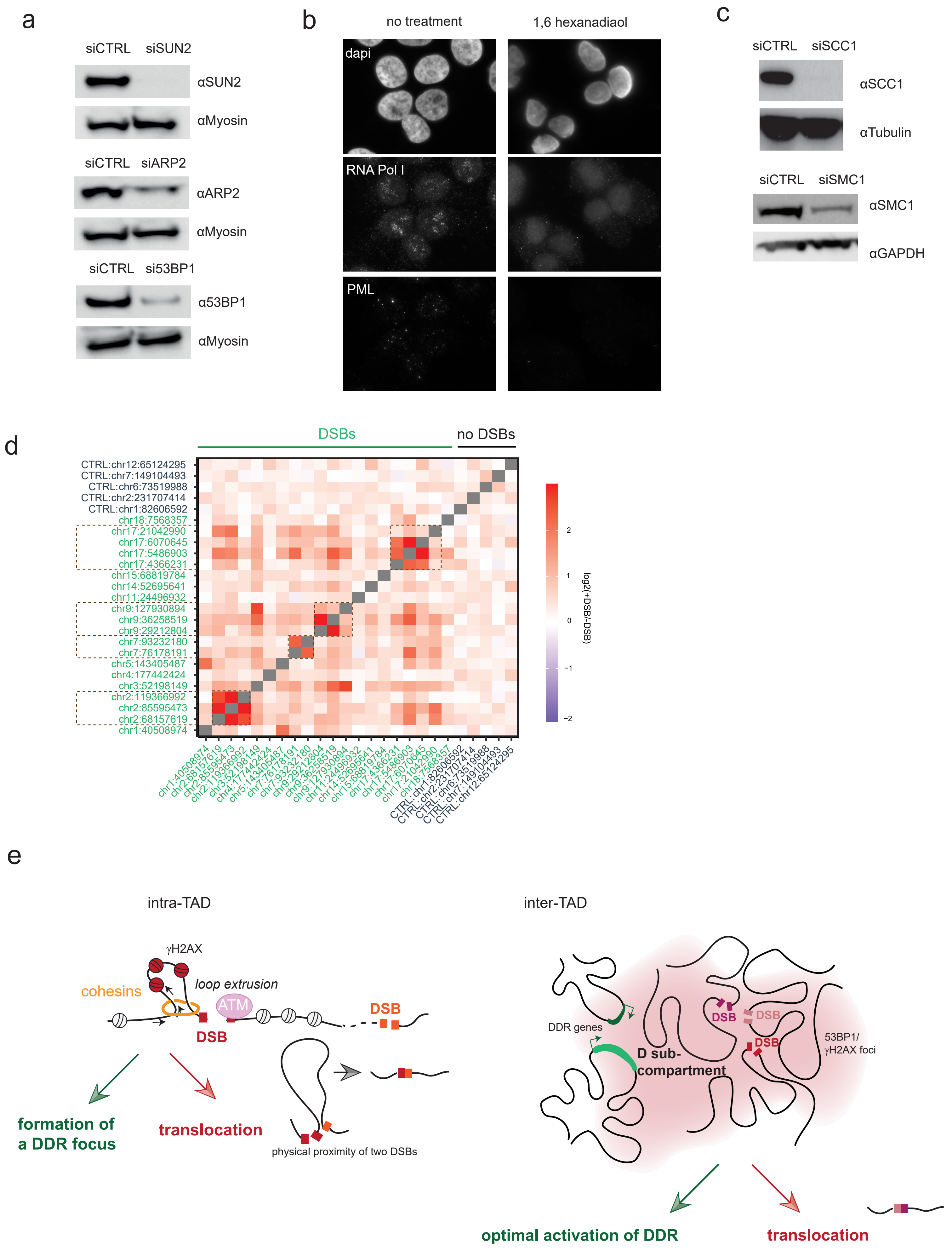
a
siCTRL siSUN2
αSUN2
αMyosin
siCTRL siARP2
αARP2
αMyosin
siCTRL si53BP1
α53BP1
αMyosin
b
no treatment
1,6 hexanadiaol
dapi
RNA Pol I
PML
c
siCTRL siSCC1
αSCC1
αTubulin
siCTRL siSMC1
αSMC1
αGAPDH
d
DSBs
no DSBs
log2(+DSB/-DSB)
e
intra-TAD
inter-TAD
γH2AX
cohesins
loop extrusion
ATM
DSB
formation of a DDR focus
translocation
physical proximity of two DSBs
DDR genes
D sub-compartment
53BP1/ γH2AX foci
optimal activation of DDR
translocation

**Figure S1. Related to Figure 1. ATM and cohesin-dependent local changes within damaged TADs**

(a) Boxplot showing the differential Hi-C read counts (log2 +DSB/-DSB) within -0.5/+0.5Mb regions containing the 80 best induced DSBs (Damaged domains, red) without inhibitors, upon ATM inhibition (ATMi) or upon DNA-PK inhibition (DNA-PKi). P-values, non-parametric wilcoxon test tested against $\mu$=0. No inhibitor *vs* ATMi or DNAPKi, P=paired wilcoxon test.

(b) same as in (a) but in cells transfected with a control (CTRL) or SCC1 siRNA. P-values, non-parametric wilcoxon test tested against $\mu$=0. siSCC1 vs siCTRL, P=paired wilcoxon test.

**Figure S2. Related to Figure 2. DSB clustering depends on ATM but not on DNAPK and cohesin, and is enhanced in G1**

(a) Hi-C contact matrix showing log2(+DSB/-DSB) on a region of the chromosome 17 at 50 kb resolution. Genomic tracks for γH2AX, 53BP1 ChIP-seq and BLESS following DSB induction are shown on the top panel. One representative experiment is shown.

(b) Same as in (A) but between a region of chromosome 17 and a region of chromosome 20.

(c) Boxplot showing the quantification of the differential Hi-C read counts (log2 (+DSB/-DSB)) between the 80 most-damaged chromatin domains in control or SCC1-depleted conditions (*cis* contacts were excluded).

(d) Boxplot showing the quantification of the differential Hi-C read counts (log2(+DSB/-DSB)) between the 80 most-damaged chromatin domains (80 DSBs) or 80 random undamaged sites (no

DSB (80 random)) in untreated cells (red), in cells treated with ATM inhibitor (green) or with DNA-PK inhibitor (grey) (*cis* contacts were excluded).

(e) Genomic tracks showing the differential 4C-seq signal (log2 (+DSB/-DSB)) at the DSB located on chr17: 61 850 870 in G1 (blue), S phase (green) and G2 (turquoise) smoothed using a 10kb span The BLESS signal after DSB induction is also shown. 4C-seq was performed using as a view point a DSB located on chr17:57 168 614. One representative experiment is shown.

**Figure S3. Related to Figure 3. DSB induction does not trigger major changes of A/B compartmentalization**

(a) Genome browser (Juicer) screenshot showing the Pearson matrix of Hi-C count on the left arm of chromosome 1, before and after DSB induction as indicated (500kb resolution). A representative experiment is shown. The bottom panel shows the A/B compartment (PC1) retrieved from two biological replicates of Hi-C experiments before (blue) and after (red) DSB induction on the same chromosomal region. The γH2AX ChIP-seq track is also shown (dark red).

(b) Saddle plots of Hi-C data obtained before DSB, after DSB, and after DSB in presence of ATMi or DNAPK inhibitor as indicated, showing interactions between pairs of 500kb bins sorted according their eigenvector value (PC1)

(c) Genomic tracks of the eigenvector (PC1) obtained from two biological replicates of Hi-C experiments before (blue) and after (red) DSB induction. The γH2AX ChIP-seq track is shown on the top (dark red) and DSB are indicated by arrows. The top panel shows an example of a DSB induced in the "A" compartment (positive PC1) which stays in the "A" compartment following

DSB induction. The middle and bottom panels show examples of two DSB which occurs in the "B" compartment (negative PC1) and switch to the A compartment following DSB.

(d) Heatmap showing the differential (log2 +DSB/-DSB) Hi-C read count for each of the 80 DSBs (1Mb window). DSBs were sorted according their Eigenvector value computed on 40kb around the DSB. DSB located in compartment A tends to cluster more.

**Figure S4. Related to Figure 3. A subset of DDR genes segregates with a DSB-induced sub-compartment**

(a) Genomic tracks of γH2AX ChIP-seq and first Chromosomal eigenvector (CEV) computed on differential (+DSB/-DSB) Hi-C matrix of chromosome 17 using a 100kb resolution (blue). Genomic regions displaying a positive CEV signal belong to the DSB-induced «D sub-compartment» (black arrows).

(b) Genomic tracks of γH2AX ChIP-seq and first Chromosomal eigenvector (CEV) computed on differential (+DSB/-DSB) Hi-C matrix on chromosome 6. Three biological replicate experiments are shown as well as the CEV obtained upon DNA-PK inhibition.

(c) Boxplot representing the quantification of the «D compartment» signal on a 1Mb window around the 80 best induced DSBs (red) and around 80 control undamaged regions (random, purple). *P*, non parametric Wilcoxon test

(d) Pearson correlation between γH2AX ChIP-seq and D compartment (positive CEV PC1 on differential Hi-C matrices) for three biological Hi-C replicates (Rep#1, Rep#2 and Rep#3) and the Hi-C performed in presence of DNAPK inhibitor as indicated.

(e) Pearson correlation score was calculated on 100kb bins between various histone modifications ChIP-seq data or RNA-seq obtained in DIvA cells and the D sub-compartment (CEV) signal computed from +DSB/-DSB (top panel)) or upon DNA-PK inhibition (+DSB+DNA-PKi/-DSB (bottom panel)) after exclusion of γH2AX covered chromatin domains.

(f) Enrichment for transcription factor motifs was analyzed for genes that displayed a positive D compartment value, as compared with genes that displayed a negative D compartment value.

(g) Box plot showing the distance between each genes upregulated following DSB induction and comprised either in the D sub-compartment or not (as indicated) and the closest DSB induced in DIvA cells.

**Figure S5. Related to Figure 4: Translocations arise from loop extrusion and D sub-compartment formation**

(a) Western Blot showing the depletion of SUN2, ARP2 or 53BP1 by siRNA.

(b) Immunofluorescence experiment showing RNA Pol I, PML and DAPI staining before and after a 1,6-Hexanediol 10% treatment as indicated.

(c) Western Blot showing the depletion of SCC1 or SMC1 by siRNA.

(d) Heatmaps showing the rate of translocations sequenced between five control undamaged regions and twenty DSB sites in the AID-DIvA cell line (expressed as the log2 (+DSB /-DSB)). n=4 independent experiments

(e) DSBs elicit profound changes in chromosome architecture that display both beneficial and detrimental outcomes. Left panel: Cohesin-dependent loop extrusion arising at DSB ensures ATM-dependent phosphorylation of H2AX on an entire TAD, allowing fast establishment of a TAD-scale DDR focus. On the other hand, DSB-anchored loop extrusion also displays the potential to bring in close proximity two DSBs located on the same chromosome, which can favor the occurrence of intra-chromosomal translocations. Right panel: Once assembled, the decorated γH2AX/53BP1 damaged TADs can further fuse together. This creates a new nuclear, DSB-induced sub-compartment, in which a subset of the DNA damage responsive genes, physically relocate, in order to achieve optimal activation. On the other hand, induced spatial proximity in D compartment also increases the frequency of translocations between DSBs and DDR responsive genes.

### Supplementary table 1: Primers sequences for Amplicon-Seq

| Name | Chr | coordinate of AsiSI site (hg19) | Primer coordinate (hg19) | sequence FW | sequence REV |
|---|---|---|---|---|---|
| **CTRL_CHR1** | chr1 | | 83072163 | GCACATGGGATTTTGCAGGAACATGTCTCTGTCACTGG | CCATCTGGCTTTAGTTCCACTTCAATCAGC |
| **CTRL_CHR12** | chr12 | | 65517948 | CTAGCATATGATAAAAATTTGGAAATAATGTGCTAGGC | GGCTATGATCATGTTTTATTTGTACTATAATTTCCAGGTC |
| **CTRL_EEF1A1** | chr6 | | 74229585 | CCTTTCTGGTATTAAACATACCTCAGCAGCC | GATCTTGGTTCATTCTCAAGCCTCAGACAGTGG |
| **CTRL_PTMAp** | chr2 | | 232572004 | CGACAGCTAAGTGCGGTGGCGATAAGGCCAGCG | CAGAGTTCCTCGGGAAAAATCATCGTCTCTGGAGCC |
| **CTRL_ZNF425** | chr7 | | 148801459 | CGCCGCAGGGAAACTCCTTCTGCCTGTCGTGGAC | CGAGTGTAACAAAAGTTTCCGCCTCAAGAGAAGCCTG |
| **SITE516** | chr1 | 40974643 | 40974513 | GCGTCCGGGAGCAGCTCCGAGGCCGCGGCG | GAAGGAGTCCCGGCCAACAGCACTGTGTTCCGA |
| **SITE709** | chr2 | 68384748 | 68384627 | CGGCAACCTGGAAACCAGACTCCAAACATTGGC | GGTCTTGGGCTCACTCCAGCCGCGCAACTCCCAG |
| **SITE716** | chr2 | 85822593 | 85822445 | TCCGGAAGCGCCATGGGCCAACGCTCGCA | CCCTCTCATTGGCTTGAACAAGTCAGGTGACC |
| **SITE722** | chr2 | 120124565 | 120124434 | GCTAGCGCCGCGGCGGGGGCTGGGCACGC | GAGCCCCCAACGGGCCTCCCGCCCGCTGCACC |
| **SITE765** | chr3 | 52232162 | 52232026 | TCCGCAGAGCCCAAGAAGTGGGCCTCTCTGC | GCCGCACCTACCGCGCGGGCCCTCACCTGCGCTG |
| **SITE871** | chr4 | 178363575 | 178363436 | TGACGACCAGGGGCAGAGGGCTGGAGCAGC | CGTGACACAACTCTCCCGCGGGCCCAGGGACG |
| **SITE919** | chr5 | 142785049 | 142784920 | CGGCGGCTCCCCCTGCTCTGACATCTTGAAGACG | CCTCTGGAGGCGGCCCCCGTAGATCGTCTCCGG |
| **SITE1025** | chr7 | 75807506 | 75807383 | GCGCCCAGGGCGAAGGTACTTAGACAGC | AGCCTCAGCCTCTGGCTCCCTCTGCAGGCAG |
| **SITE1031** | chr7 | 92861490 | 92861341 | TCACACTCCTAGGGGACATCGCGTCAATCTGGC | GCTGGGAACTGTAGTCTTCTGGACGCCGGCGGTAGCAACG |
| **SITE1123** | chr9 | 29212799 | 29212659 | TCAACCTCCACAGAGAACCCAGTCCGAAATCG | GCCTCAAGGTAAGAGGCAGCAAAGCTGCTGCTTG |
| **SITE1129** | chr9 | 36258513 | 36258383 | GCCACGAAGCAGGCAGAGCGCGAGC | CATGGAAGATGGTCCGCTGGTCAGC |
| **SITE1159** | chr9 | 130693170 | 130693031 | TCGCCCCCGCAGCAGCTTCCCGGGCTTTGGCCGC | CAGGCGGCCCTAGCGACCCGAGTCCCCACGCCG |
| **SITE72** | chr11 | 24518475 | 24518337 | CAATTATCGGCAGATTAGGTTTCTACTCTCCGTG | CAGGAGTGATGTCACCGGCGAGGCGAGCCCTGCC |
| **SITE219** | chr14 | 54955825 | 53162224 | CACGCTTGGGAGGCCAGTCCGCACGCTCGGTCG | GCGGGGCCGAGCCGGGAAGCTGGTCGGTGCG |
| **SITE270** | chr15 | 69112120 | 69111977 | CTGCCGACAAGCCCAGCACTGAGAAAGACGGGC | TCCACCCCACCTTCGGGCCAGATGCTGATGTTTCT |
| **SITE341** | chr17 | 4269523 | 4269398 | CGGGCGAGAACGGCTGGGCCCGGCCGGGACCG | AGCCCCGGCGCCAGCCCGGGCCCTCGGC |
| **SITE343** | chr17 | 5390220 | 5390088 | CGGCCTGTGGGTCGGCCTCACCCCGGCCTCCG | CCGCCTCATCTTCTCTGCTAGACTTAGAGTTCCTG |
| **SITE344** | chr17 | 5973962 | 5973831 | GTCCGCTTGCCGCGGGCGGCCGGAGACGTGC | CGCCGCGCTCCCAACCGCGTCTGCAC |
| **SITE360** | chr17 | 20946300 | 20946179 | CGGCGGCGAGGGAGGCGAGGACGACGCCAG | GTGACGTAATCTCCGTCCGCGGCCGCGGCG |
| **SITE396** | chr18 | 7566712 | 7568200 | GAGTCCCTGGCCGGCTGCAAAAGGAACAGCAG | AGCCTCTCCGCAGACGCTGCACGACCGCG |

#### Supplementary table 2: Primer sequences for 4C-seq

| Name | Forward primer | Reverse primer |
|---|---|---|
| Viewpoint DSB1 (chr1, cluster-prone) | AATGATACGGCGACCACCGAGATCTACACTCTTTCCCTACACGACGCTCTTCCGATCTAACCTGGCAACTTATGAATCAGGA | CAAGCAGAAGACGGCATACGAGATNNNNNNGTGACTGGAGTTCAGACGTGTGCTCTTCCGATCTATGTCAAAAGCCAAGGGGACA |
| Viewpoint DSB2 (chr17, cluster-prone) | AATGATACGGCGACCACCGAGATCTACACTCTTTCCCTACACGACGCTCTTCCGATCTTCCTTACGATTATTTGTGAATTTTG | CAAGCAGAAGACGGCATACGAGATNNNNNNGTGACTGGAGTTCAGACGTGTGCTCTTCCGATCTAAGCTAATTCTGAGTTACATACATT |
| Viewpoint DSB3 (chr21, not cluster-prone) | AATGATACGGCGACCACCGAGATCTACACTCTTTCCCTACACGACGCTCTTCCGATCTGATTACGTAGAAGGGTGCC | CAAGCAGAAGACGGCATACGAGATNNNNNNGTGACTGGAGTTCAGACGTGTGCTCTTCCGATCTAAGGCAAATGATAACCCTGT |
| Viewpoint DSB4 (chr20, cluster-prone) | AATGATACGGCGACCACCGAGATCTACACTCTTTCCCTACACGACGCTCTTCCGATCTGGTTATACTAAGATGTCAGTTCCT | CAAGCAGAAGACGGCATACGAGATNNNNNNGTGACTGGAGTTCAGACGTGTGCTCTTCCGATCTCACGCACCTGGTTTAGATT |
| Viewpoint ctrl region (chr17) | AATGATACGGCGACCACCGAGATCTACACTCTTTCCCTACACGACGCTCTTCCGATCTTCCTCAGGTTATCATCCCAA | CAAGCAGAAGACGGCATACGAGATNNNNNNGTGACTGGAGTTCAGACGTGTGCTCTTCCGATCTCACCTTCGCTGTACCTTTG |

NNN is the position of the optional index
